# Mitochondrial Response to Psychological Stress and Its Medial Prefrontal Biomarker Correlates

**DOI:** 10.1101/2025.07.30.667787

**Authors:** A. Ankeeta, Ashutosh Tripathi, Yizhou Ma, Bindu Pillai, Joshua J. Chiappelli, Jennifer N. Jernberg, Alia Warner, Keiko Kunitoki, Bhim M. Adhikari, Si Gao, Xiaoming Du, Loise Kabui, Francisco Pallares Solano, Oluwabunmi Akindona, Zhenyao Ye, Shuo Chen, Mohammad Milad, Peter Kochunov, Anilkumar Pillai, L Elliot Hong

## Abstract

**Background:** Stress response obligates increased mitochondrial activities to meet stress-induced high energy requirement. This stress–mitochondrial response process involves glucocorticoid but also multiple alternative pathways that are top-down regulated by the medial prefrontal cortex (mPFC). These pathways are important for many neuropsychiatric conditions that are sensitive to stress. However, the field lacks a reliable, clinically accessible stress–mitochondrial response paradigm to study the process in humans.

**Method:** We used an established psychological stress challenge combined with assaying salivary cell-free mitochondrial DNA (cf-mtDNA), thought to reflect heightened mitochondrial changes or disruptions, in 35 healthy individuals (21 males). We also explored if these stress-induced cf-mtDNA marker elevations were associated brain metabolites as measured by magnetic resonance spectroscopy (MRS), as well as high-resolution brain imaging based cortical thickness focusing on the mPFC.

**Results:** We found that salivary cf-mtDNA was significant elevated immediately after the stress challenge (p=2.0×10^-7^) and gradually declined after. Exploratory causal analysis showed that this cf-mtDNA response was not primarily driven by cortisol response. Instead, individuals with higher baseline dACC lactate+ levels, thought to in part reflect mitochondrial dysfunctions, was significantly associated with the cf-mtDNA response (r=0.80, p<0.001). Higher mtDNA response was also significantly associated with thinner dorsomedial prefrontal cortex (r=-0.52, p=0.01). Age had a U-shape effect such that cf-mtDNA response trended lower in earlier adulthood but higher in older people, explaining 33.8% of the ct-mtDNA response variance (p=0.003).

**Conclusion:** This stress challenge-salivary cf-mtDNA assay paradigm may offer a new, non-invasive approach to evaluate the stress-mitochondrial pathway functioning in aging, psychopharmacology, and neuropsychiatric conditions where psychological stress plays a role.

## Introduction

Stress plays a prominent role in health and major psychiatric disorders such as depression, schizophrenia and many others. Stress response is a highly energetic process involving activation of the hypothalamic-pituitary-adrenal (HPA) axis and release of glucocorticoids that mobilize energy metabolism through increasing mitochondrial activity [1, 2]. Mitochondria also undergo adaptive response by interacting with catecholamine, neuroendocrine, metabolic, inflammatory, and transcriptional pathways during psychological stress [3, 4]. Mitochondria contain circular, double-stranded mitochondrial DNA (mtDNA) and are central to cellular stress response through energy production in the form of ATP. Both acute [5–7] and chronic stress [3, 8] conditions have been reported to alter mitochondrial function and increase in the release of mtDNA into circulation. We have shown that chronic stress induces increase plasma cell-free mtDNA (cf-mtDNA) in animals that is causally linked to social behavioral deficits, suggesting that cf-mtDNA can serve as a functional biomarker for the stress-mitochondrial response pathway [9]. Whether this model is translatable to humans is unclear. In this study, we will address whether the stress-mitochondrial pathway can be similarly assayed in humans.

In addition to its presence in blood, cf-mtDNA may also be released into saliva as a result of cellular stress and damage [10]. Salivary cf-mtDNA has recently been shown to be feasible to obtain [11–13], which were modestly correlated with the diurnal cortisol levels [11, 14]. We are aware of three studies attempting to elicit cf-mtDNA responses using psychosocial stressors based on the Trier social stress test (TSST) paradigm. A study of N=20 healthy people showed that TSST induced significant salivary cf-mtDNA response within 15 minutes, although it also showed significant increases cell-free nuclear DNA (cf-nDNA) yet the study did not correct for the cf-nDNA, raising a question whether the finding was cf-mtDNA specific [5]. A larger study of N=46 healthy participants found that TSST failed to elicit a significant salivary cf-mtDNA response [7]. The same study reported that the plasma cf-mtDNA was significantly increased but it was peaked only at 45 minutes after TSST (see Supplementary Figure 5B of the study), raising a question as to the nature of this response. The latest effort using also the TSST paradigm included N=72 healthy participants, and concluded that their version of the task also did not significantly elicit TSST-induced cf-mtDNA in either serum or plasma [13]. Interestingly, salivary cf-mtDNA was found not significantly responded to the psychosocial stress paradigm despite a robust and significant salivary cortisol response [7], further suggesting that a glucocorticoid response may not always translate to a cf-mtDNA response based on the paradigm. However, we are unaware of report that has explicitly tested this glucocorticoid – cf-mtDNA response causality in humans. Thus, we used another psychological stress paradigm to potentially elicit salivary mtDNA response and model its relationship with cortisol response.

Besides glucocorticoids, stress response is heterogeneous across people [15], driven by central regulations based on individual brain structural and functional capacities. In rodents, the medial prefrontal cortex (mPFC) is the center for top-down emotional, motivation, and fear regulations in connections with the hippocampus and amygdala [16, 17]. The human equivalent of the mPFC includes the dorsomedial prefrontal cortex (dmPFC) and dorsal anterior cingulate cortex (dACC) that are part of the “central autonomic–interoceptive network” that respond to heightened arousal states during fear conditioning and other stressful events [18]. Impairment in mPFC increases the vulnerability to stress-related disorders and abnormal response to stress [19]. It is currently unknown if mitochondrial response to stress may be regulated by the mPFC hub. Therefore, the dmPFC and the dACC area was selected based on its role in top-down regulation of the stress response and emotional appraisal [20, 21]. Specifically, we included prefrontal cortical thickness as a measure of the prefrontal structural integrity, and examined its relationship with cf-mtDNA response to stress. We focused on cortical thickness (a structural marker) rather than functional data because structural measures offer a stable, trait-like index of brain integrity, less influenced by transient states or task demands [20–22], and impairment of mPFC structure has been consistently linked to stress vulnerability in both animal and human studies [21, 22].

While mitochondrial function in the human brain can be assessed in vivo using PET imaging with tracers targeting mitochondrial complex I (e.g., [18F]BCPP-EF;) [23], this technique is not yet widely accessible in clinical or research settings due to the isotope requirement and the invasive nature. Therefore, as a feasible proxy, we used brain metabolites measured via magnetic resonance spectroscopy (MRS), to test whether psychological stress-induced cf-mtDNA may be linked to brain metabolites. Our specific interest is one of the metabolites lactate as it has been mechanistically linked to mitochondrial metabolisms. Mitochondria are primarily energized by pyruvate, which is generated from glucose or lactate [24]. Lactate is traditionally considered as a anaerobic metabolite of glucose metabolism, but modern evidence showed that lactate is directly oxidized by mitochondria especially during higher energy demand and serves as a signaling molecule [24–27] critical for maintaining brain energy homeostasis and supporting neuronal function [28, 29]. High lactate can damage mitochondria and cause mtDNA leakage [30]. We have shown that greater stressor exposure was associated with higher brain lactate+ and N-acetylaspartate (NAA) levels in schizophrenia patients [31], linking stress to these bioenergetics metabolites. Therefore, we used a magnetic resonance spectroscopy (MRS) to broadly explore several metabolites at the dACC [32, 33] to test the hypothesis that bioenergetics metabolite levels may index central mitochondrial vulnerability and thus may be associated with higher cf-mtDNA response to stress.

In summary, we tested four related stress-mitochondrial pathway hypotheses in healthy humans: 1) whether the stress-mitochondrial pathway can be assayed using salivary cf-mtDNA using a psychological stress challenge, 2) whether the response is glucocorticoid-dependent, 3) whether bioenergetic-related metabolites in the dACC may be associated with the vulnerability to have higher stress-induced cf-mtDNA, and 4) whether this stress-induced cf-mtDNA response is in part determined by prefrontal integrity as measured by dmPFC cortical thickness as compared to other parts of the cortex.

## Methods and Materials

### Participants

The study included 35 healthy participants, average age 43.03±15.62 years, 21 males. Structured Clinical Interview for DSM-4 or 5 was used to exclude major psychiatric diagnoses including schizophrenia spectrum disorders, bipolar disorder, major depressive disorder, anxiety disorders, or other conditions that could significantly affect cognitive or emotional functioning. Other exclusion criteria included a history of neurological conditions, head trauma with cognitive sequelae, and active and uncontrolled major medical conditions. Participants with recent substance abuse or dependence other than nicotine or cannabis in the prior 6 months were excluded, no participants met criteria for current cannabis use disorder. Demographic and clinical measures are in **Table 1**. Data were collected under protocols approved by the University of Maryland Institutional Review Board. Participants provided written informed consent.

**Table 1.**
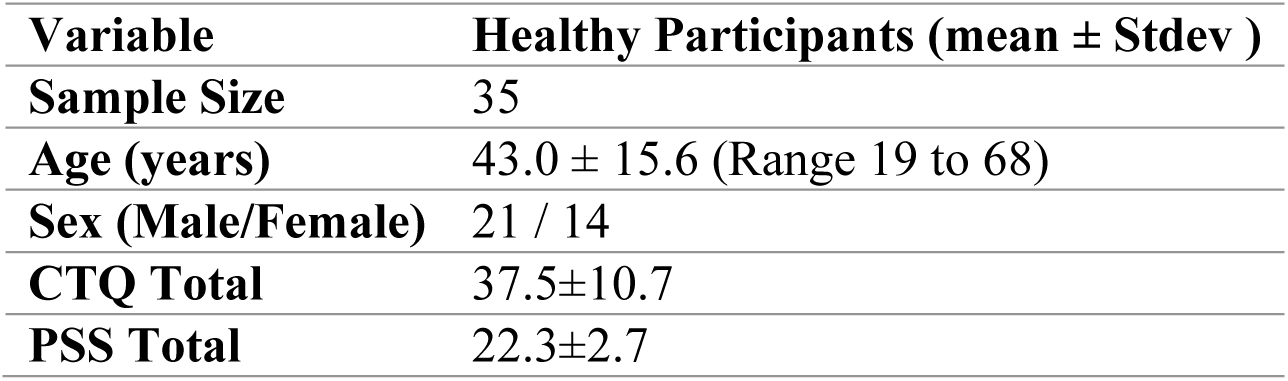
Demographic and Clinical Characteristics of the Participants.

| **Variable** | **Healthy Participants (mean ± Stdev )** |
| --- | --- |
| **Sample Size** | 35 |
| **Age (years)** | 43.0 ± 15.6 (Range 19 to 68) |
| **Sex (Male/Female)** | 21 / 14 |
| **CTQ Total** | 37.5±10.7 |
| **PSS Total** | 22.3±2.7 |

### Psychological stress challenge

Participants completed two psychological distress-inducing tasks, the order of which varied randomly. One was the computerized Paced Auditory Serial Addition Task (PASAT) by using a computer mouse to perform rapid arithmetic additions [34]. Incorrect response or failed to respond resulted in a loud (∼90 decibel) aversive explosion sound. Individual speed of response and accuracy were titrated to provide similar challenges across participants. Another was the Mirror-Tracing Persistence Task (MTPT) [35] by tracing a dot along a star on the screen using the mouse. Tracing was challenging because the cursor movement was opposite to that of the mouse. Tracing outside the line or keeping stationary was met with the explosion sound. The width of the star outline was titrated depending on performance. Participants could quit, but were informed that the better they performed the greater the monetary bonus to provide strong motivation to persist. Although incentives were modest, they have been effective in prior DI studies using similar paradigms. Both tasks assess distress intolerance (DI) [36–40], thought to capture a person’s inability to tolerate distress [39]. We combined both tasks to reduce biases due to cognitive vs. manual skill. After the test, participants quietly watched a natural scenery video. Salivary samples were collected at baseline, immediately after (time 0) or if quitting, 20-, and 40-minutes post-stress, and stored in -80°C. DI was defined as those who quit both tasks, otherwise non-DI.

### Salivary cf-mtDNA and cortisol assays

Total DNA was isolated from 200μl saliva using DNA isolation/purification kits (Qiagen, Hilden, Germany) according to the manufacturer’s protocol. Quantity of mtDNA was measured by qRT-PCR by analyzing the mitochondrially encoded NADH dehydrogenase subunit 1 (ND1) gene (forward primer-5’ ATACCCATGGCCAACCTCCT 3’ and reverse primer-5’ GGGCCTTTGCGTAGTTGTAT 3’). Primers were synthesized by Integrated DNA Technologies. MasterCycler (Eppendorf, NY) to perform qRT-PCR using iTaqTM Universal SYBR® Green Supermix (Bio-Rad, CA). Mitochondrial DNA levels were adjusted for nuclear DNA (nDNA) levels using the average values of three nuclear genes; 18 S rRNA (forward primer-5’ AGAAACGGCTACCACATCCA 3’ and reverse primer-5’ CCCTCCAATGGATCCTCGTT 3’), B2M (forward primer-5’ CACTGAAAAAGATGAGTATGCC 3’ and reverse primer-5’ AACATTCCCTGACAATCCC 3’), and β-actin (forward primer-5’ GGACTTCGAGCAAGAGATGG 3’ and reverse primer-5’ AGCACTGTGTTGGCGTACAG 3’). Analysis of mtDNA used the 2^delta delta Ct method [41, 42]: cycle threshold (Ct) values were collected after qRT-PCR. Delta Ct of each sample was calculated by subtracting the Ct values of mtDNA from the nDNA. Then delta delta Ct was calculated by subtracting delta Ct of the post-stress time points from baseline delta Ct. Relative quantity (RQ) of mtDNA was calculated using delta delta Ct[9, 43, 44].

Cortisol assay followed previously reported methods [45, 46]. Briefly, samples were thawed and centrifuged at 10,000 g for 10-min. Cortisol was assayed using a commercial enzyme immunoassay kit (Salimetrics) following the manufacturer’s protocol. Intra-assay coefficient of variance (CV) was 7.77% and inter-assay CV was 4.18% in our lab [47].

The overall response was evaluated using the Area Under the Curve (AUC) for cf-mtDNA RQ and cortisol across the 4 time points calculated by the trapezoidal rule:

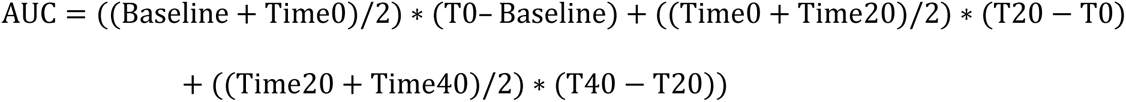

### Structural imaging and MRS

Proton magnetic resonance spectroscopy (¹H-MRS) and structural imaging acquisitions were conducted using on a Siemens MAGNETOM 3T scanner with a 64-channel head coil and a high-performance gradient system. T1-weighted images were acquired using a fast spoiled gradient recalled sequence (TR=2400ms, TE=2.22ms, flip angle=8°, matrix=300×320, slices per slab=208) at 0.80mm isotropic. Imaging processing used FreeSurfer’s recon-all pipeline [48], which includes skull stripping, surface reconstruction, and quality control on cortical surfaces, subcortical segmentation, and image alignment. Cortical thickness was extracted using the Desikan-Killiany atlas [49] preinstalled in FreeSurfer for 34 regions per hemisphere. MRS spectra were acquired from a 40×30×20mm voxel centered on the dACC using PRESS (TR/TE=2000/30ms), 2000 Hz spectral width, flip angle=90°, 2048 complex points, auto phase cycling, NEX=128. A water suppressed spectra (NEX=16) was acquired for water scaling, eddy current correction, and quantification. We used this standard single-voxel spin echo sequence to obtain a broad neurochemical profile survey across multiple metabolites, but the short TE (30 ms) does not separate metabolites like lactate from lipid and other contaminants as compared to longer TEs (e.g., 144 ms) or editing sequences (e.g., MEGA-PRESS) that are typically needed to increase lactate specificity from lipid and threonine [50, 51]. Meanwhile, the current sequence retains high signal-to-noise ratio across a wider metabolic range, and the lactate identified at 1.33 ppm by a characteristic doublet can be estimated and labeled as lactate+ (lac+). Similar approaches using TE around 30 ms have successfully quantified lactate+ in clinical populations, including schizophrenia and bipolar disorder [52–57]. LCModel (http://s-provencher.com/lcmodel.shtml) was used for quantification using a simulated basis set of 19 metabolites, with lactate+ as the a priori metabolite but other metabolites were also reported to evaluate specificity. Metabolite levels were referenced to water and reported in institutional units. Spectra with FWHM>0.1 ppm and signal to noise ratio<10 were excluded. We used Cramér-Rao lower bounds (CRLB) %SD≤20% for most of the metabolites, but %SD≤30% for lactate+, GABA and glutamine, as the 20% threshold may be too conservative for low concentration metabolites [31, 58]. Parameters exceeding these criteria were considered outliers. The proportion of the cerebrospinal fluid (CSF) and white matter within the voxels was calculated based on segmentation of MP-RAGE images. Metabolite values were corrected by dividing raw values by (1-CSF fraction) [59]. Examples of voxel placement and spectra are shown in **Figure 2A&B**. All metabolites listed (Ins, NAA, Cho, PCr, Glu+Gln, Cr, GSH, Asp, sI, Tau, and NAAG) had a %SD below the 20% threshold. However, Lactate+ (%SD=24%) and GABA (%SD=30%) were slightly higher but were retained [60, 61]. Final metabolites included were based on quality criteria and biological relevance.

### Clinical and cognitive assessment

The Childhood Trauma Questionnaire-Short Form (CTQ) was used to assess developmental stressful experiences consisting of physical, emotional, and sexual abuse, and neglect during childhood [62]. The Perceived Stress Scale (PSS) was used to measure subjective levels of stress over the past month [63]. Cognitive testing included the digit symbol coding task to assess information processing speed and the digit sequence task to assess working memory.

### Data analysis

All analyses were conducted using IBM SPSS Statistics version 28. To evaluate changes in salivary cf-mtDNA and cortisol across four time points, we used repeated measures ANOVA, applying Greenhouse-Geisser correction when sphericity assumptions were not met. To explore the moderating effect of distress intolerance (DI), DI was added as a between-subjects factor in the model. Associations among cf-mtDNA, cortisol, and clinical variables were examined using Pearson’s correlation and multiple linear regression, controlling for age and sex. Given the number of comparisons, False Discovery Rate (FDR) correction was applied to metabolite analyses to reduce the likelihood of false positives. For other analyses, results were considered significant at p < 0.05 (two-tailed), with cautious interpretation due to the small sample size.

We tested an exploratory mediation model examining whether dACC lactate+ mediated the association between cortical thickness and stress response, based on prior evidence linking neuroenergetics to brain structure and stress physiology and given the pilot nature of the study, these analyses were considered hypothesis-generating. Taking advantage of the time course data, we further investigated causal relationships among cf-mtDNA and cortisol responses using the constraint-based causal discovery algorithm [64], to test whether the cf-mtDNA response was driven by glucocorticoid response or vice versa, or they were independent. Briefly, we tested conditional independence by conducting two partial correlation analyses: i) regressing cortisol on stress while controlling for mtDNA and ii) regressing mtDNA on stress while controlling for cortisol. If two variables become independent when conditioned on the third, we eliminated the edge between them (**Figure 1F**). We then oriented the remaining edges using orientation propagation rules [64, 65]. Specifically, if a variable was directly linked to one variable but not another: stress→mtDNA – cortisol or stress→cortisol - mtDNA), we treated the conditional variable as a collider denoted as △ 𝒁 and inferred the directionality [64]. To explore potential non-linear age-related effects, we included a quadratic term (age) in models assessing associations between biological variables. This was based on evidence that structural and metabolic brain markers may follow non-linear trajectories over the lifespan [66, 67].

**Figure 1.**
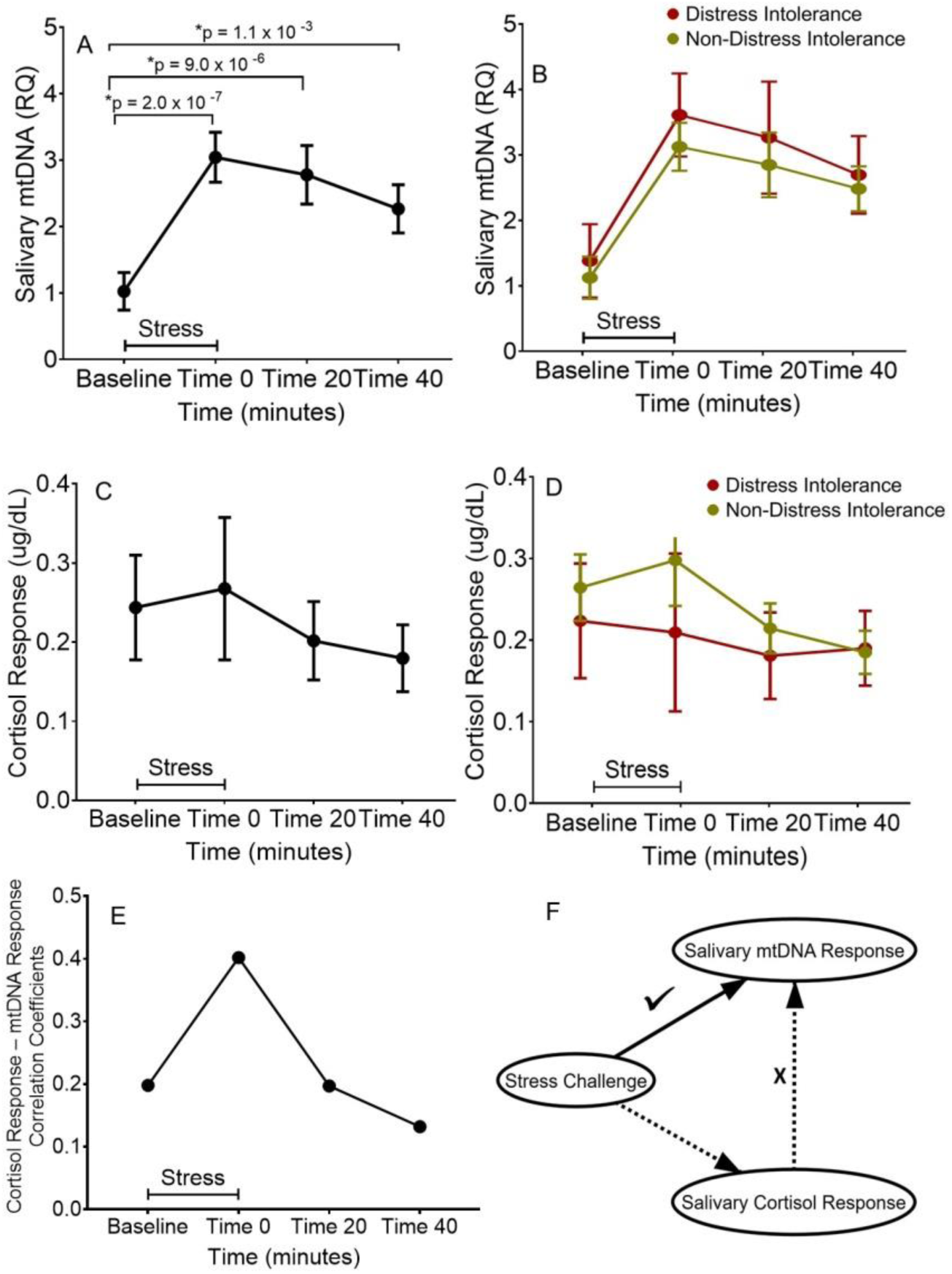
**A**: Mean (and standard error) salivary cf-mtDNA levels at baseline and time 0, 20, and 40 minutes post stress. **B**: Similar trends were found when the sample was divided by distress intolerance. **C** and **D**: Salivary cortisol levels over time. **E**: Correlations between cf-mtDNA and cortisol at each time point, showing increased correlation immediately after the stress. **F**: Discovery causal modeling failed to show a significant stress→cortisol→cf-mtDNA response pathway, where there was a robust stress→cf-mtDNA response pathway independent of the cortisol response.

## Results

In all analyses, cf-mtDNA response was quantified using relative quantification (RQ) normalized to baseline values. We assessed all discrete timepoint changes (e.g., 20 minutes post-stress vs. baseline), including area under the curve (AUC) as overall indicators of stress response. Cortisol response was analyzed using both individual time points and AUC. Distress Intolerance (DI) was modeled as a between-subject variable to test moderation effects on within-subject responses over time.

### Cf-mtDNA response to psychological stress

The sample includes 35 participants with salivary cf-mtDNA RQ data available. Among them 5 have one to two missing data points, which were imputed using the Expectation-Maximization (EM) algorithm. EM was chosen to preserve sample size in repeated-measures ANOVA. To validate robustness, we additionally conducted linear mixed-effects models (LMMs), which do not require imputation and handle unbalanced designs. Results were consistent across approaches. Missingness was primarily due to low saliva yield or delayed collection timing; no imputation was done for missing imaging data. In the LMM, pairwise comparisons indicated that Time 1 (Baseline) significantly differed from Time 2 (0 minutes, p<0.001), Time 3 (20 minutes, p<0.001), and Time 4 (40 minutes, p=0.009), with lower cf-mtDNA RQ values at Time 1 compared to subsequent time points. No other contrasts were significant after correction for multiple comparisons. Consistent with these findings, repeated measure ANOVA also showed that there was a significant effect of time (F(1,34)=16.37, p<0.001) with Time 1 significantly differed from Time 2 (p<0.001), Time 3 (p<0.001), and Time 4 (p=0.001), peaked immediate after the stress and returned about half way towards baseline by 40 minutes (**Figure 1A**). The time×DI interaction was non-significant (p=0.95), suggesting that DI was not a major factor for the cf-mtDNA response (**Figure 1B**).

### Cortisol response to stress

There was a significant effect of time (F(1,33)=6.0, p=0.04) although post-hoc tests found no significant increases in any post-stress time point compared to baseline (**Figure 1C**). The time×DI interaction was also non-significant (p=0.68) **(Figure 1D).**

### Relationship between cortisol and mtDNA response to stress

Across the 4 time points, there was a significant correlation between cortisol and mtDNA responses only at immediately after the stress test (r=0.36, p=0.04) (**Figure 1E**), suggesting a potential association in their responses. A causal discovery modeling **(Figure 1F)** showed that there was a possible causal relationship between cortisol and cf-mtDNA response at the first post-challenge time point. The causal direction identified was cortisol→mtDNA (Estimate=2.16, SE=1.66, t=1.31); however, this effect was not statistically significant (p=0.20) controlling for age, sex and DI. This suggests that while cortisol may influence cf-mtDNA changes, the relationship was not confirmed as a direct causal pathway, while stress and mtDNA remained significantly connected **(Figure 1F)**.

### Relationship between brain metabolites and mtDNA response to stress

Giving the lack of evidence of a primarily glucocorticoid mechanism, we tested alternative brain mechanisms (**Figure 2**). Sixteen participants have both MRS and salivary mtDNA data were available. Using the salivary mtDNA RQ AUC as the overall response measure, only lactate+ showed a strong positive correlation (r=0.80, p<0.001), significant after correction for multiple comparisons, (**Figure 2C-D**), indicating that individuals with high baseline dACC lactate+ had significantly higher mtDNA response to stress. No other metabolites showed significant effect after correction for multiple comparisons (**Figure 2C**), although this was based on a small sample and confirmatory studies would be needed.

**Figure 2.**
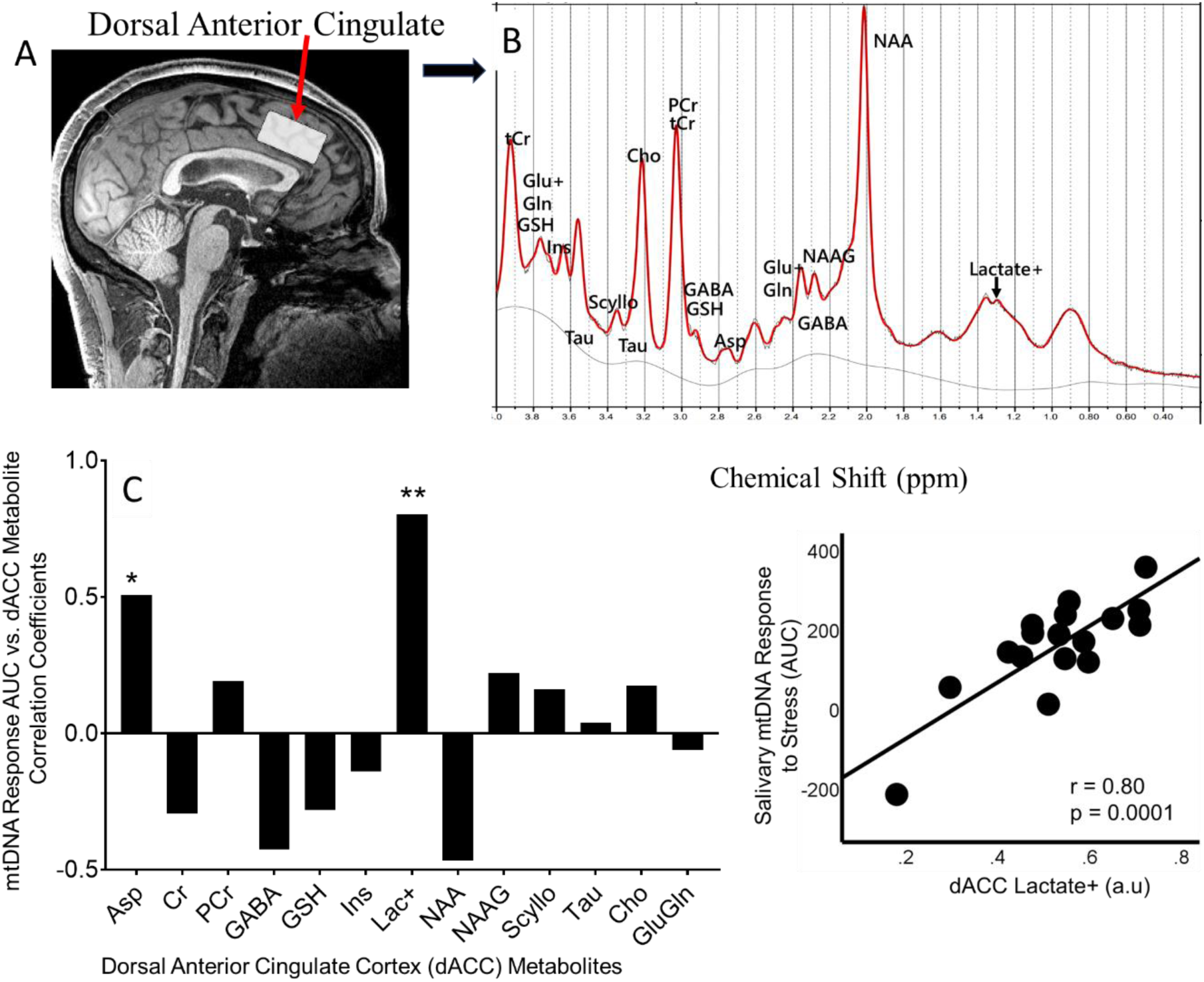
Spectroscopic analysis at dorsal anterior cingulate cortex (dACC) metabolites and their associations with salivary cf-mtDNA response. **A**: Voxel placement. **B**: A representative metabolite quantification output from LCModel. **C**: Correlation coefficients between metabolites measured by ^1^H-MRS at dACC and salivary cf-mtDNA response measured by the area under the curve (AUC) of the relative quotients (RQ), with the strongest and the only statistically significant positive correlation observed at lactate+ (r=0.80, p<0.001) (**D**). Metabolites listed are Aspartate (Asp), Creatine (Cr), Phosphocreatine (PCr), Gamma-aminobutyric acid (GABA), Glutathione (GSH), Myo-Inositol (Ins), Lactate+ (Lac+), N-Acetylaspartate (NAA), N-Acetylaspartylglutamate (NAAG), Scyllo-inositol (Scyllo), Taurine (Tau), Choline (Cho), Glutamate (Glu), and Glutamine (Gln). a.u.: arbitrary unit. * nominally significant; ** significant after corrections for multiple comparisons.

### Relationship between dmPFC cortical thickness and mtDNA response to stress

Cortical thickness data was available in thirty participants. The correlation analysis with mtDNA response (AUC) revealed negative associations in multiple frontal regions (**Figure 3A**) with the strongest association at left dmPFC along the superior frontal lobe (r=-0.52, p=0.003; p=0.01 adjusted using the FDR/Benjamini-Hochberg method) (**Figure 3B**), such that thinner cortex in this area was associated with a higher mitochondrial response to stress. Additionally, the left frontal pole (r=-0.39, p=0.03) and the right rostral middle frontal cortex also demonstrated nominally significant correlations (r=-0.40, p=0.026); these were not significant after FDR correction.

**Figure 3.**
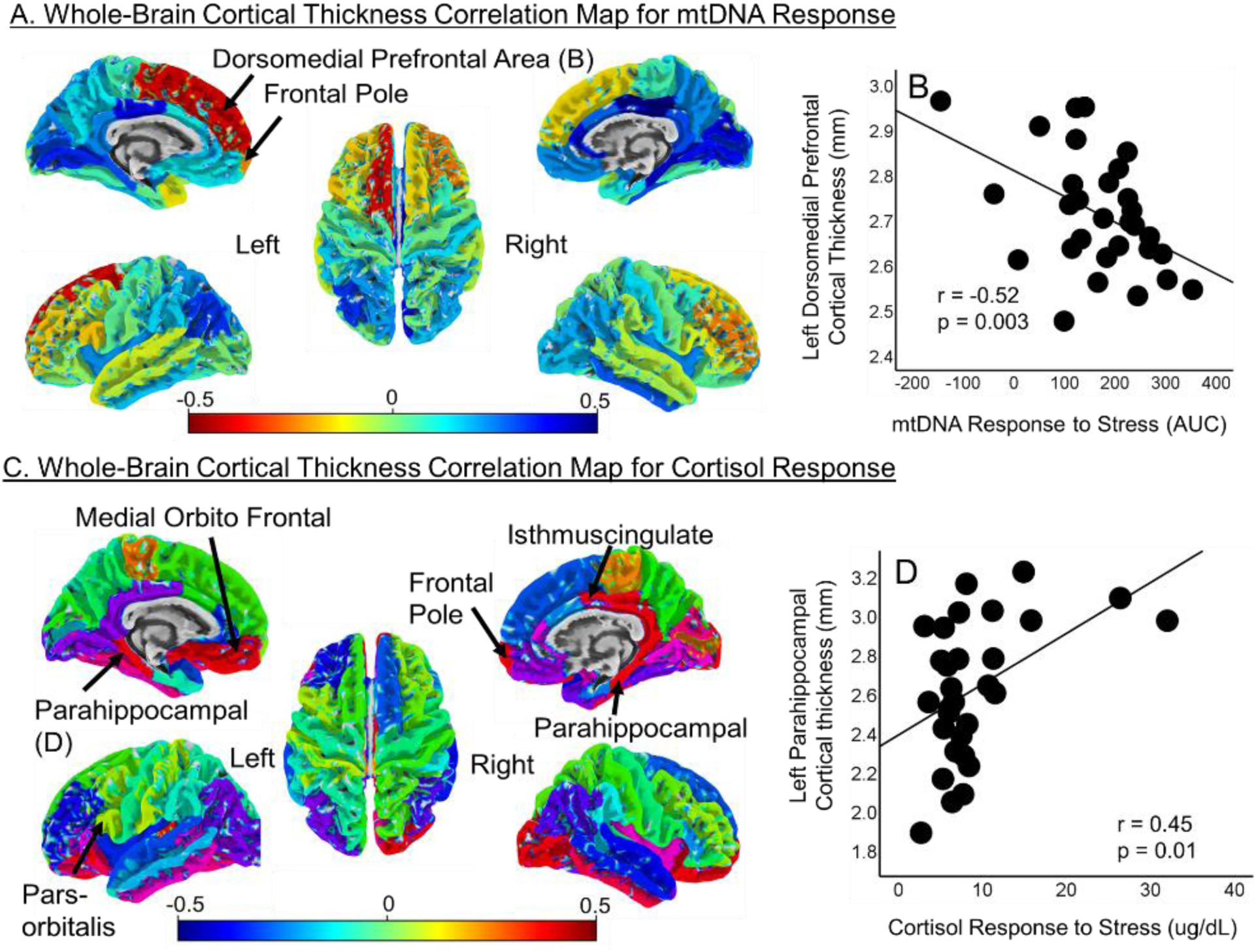
**A**: Whole-brain cortical thickness analysis identifying regions significantly associated with salivary cf-mtDNA response area under the curve (AUC). **B**: Left dorsomedial prefrontal cortex (dmPFC) area showed the strongest and negative association with cf-mtDNA response. **C**: Parallel analysis for salivary cortisol response suggested a different pattern of correlations with cortical thickness. **D**: Left parahippocampal gyrus area showed the strongest and positive association with cortisol response.

For specificity testing, we also plotted the correlation coefficients with cortisol response (AUC), and found positive associations in several regions, with the left parahippocampal cortex exhibited the strongest correlation (r=0.46, p=0.01) (**Figure 3C-D**). The left medial orbitofrontal, left pars orbitalis, right isthmus cingulate, right parahippocampal cortex, and right frontal pole (r=0.36 to 0.43, p=0.049 to 0.017) also demonstrated nominally significant correlations; none was significant after corrected for multiple comparisons; but a key observation is the different cortical association patterns between cf-mtDNA and cortisol response to stress.

### High lactate+ may affect dmPFC effect on cf-mtDNA response

To address this, we first plotted the associations between dACC lactate+ level and whole-brain cortical thickness (**Figure 4A**), and found that high lactate+ was most strongly associated with left dmPFC cortical thickness (**Figure 4B**). Using lactate+ as the mediator, the association between dmPFC and cf-mtDNA AUC was reduced but still significant (p=0.02), while there was a significant indirect effect through lactate+ (p=0.016) (**Figure 4C**), supporting that the dmPFC effect on cf-mtDNA response to psychological stress was partially but significantly mediated by dACC lactate+ level.

**Figure 4.**
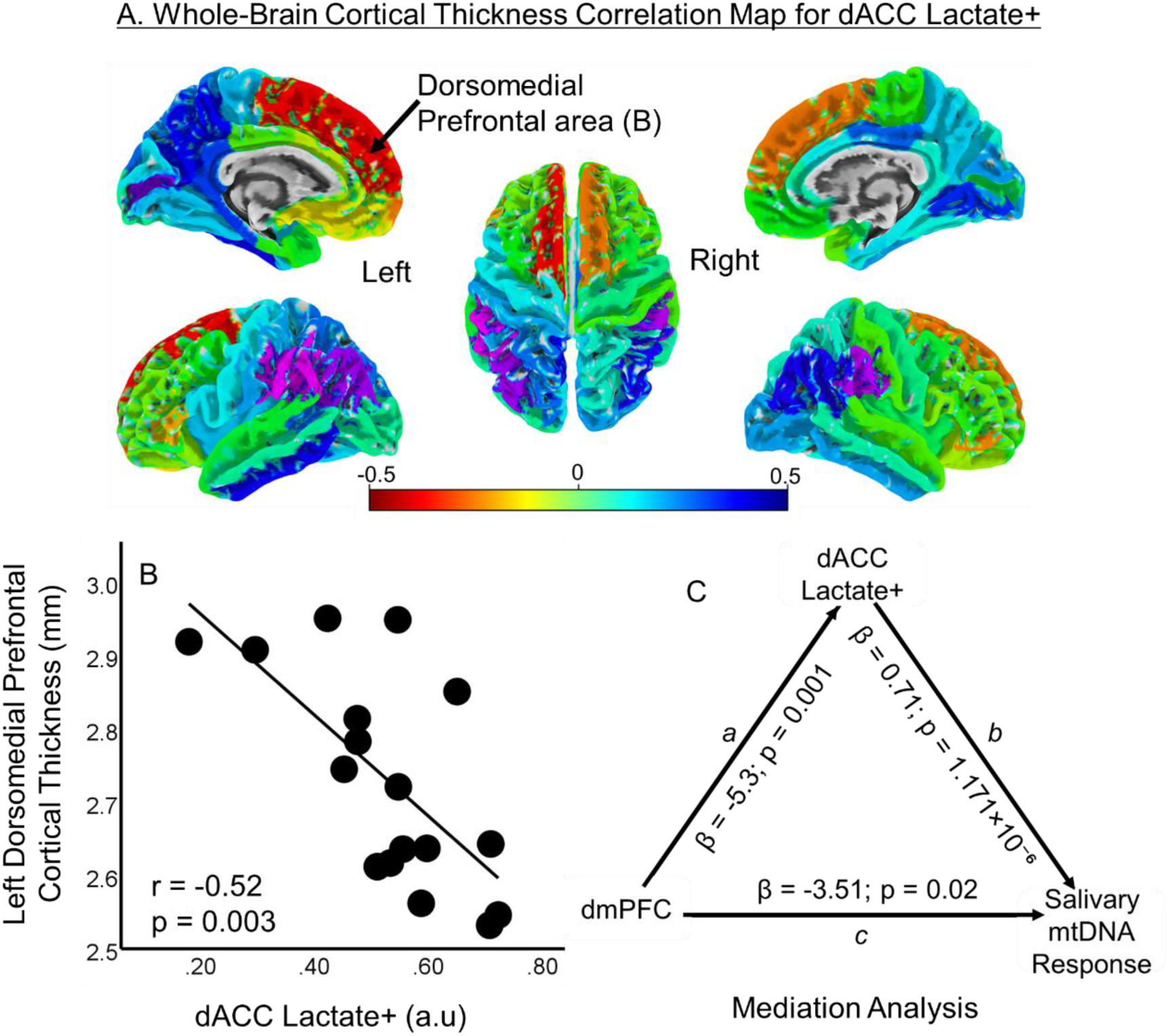
**A**: Lactate+ measured at dorsal anterior cingulate cortex (dACC) was associated with cortical thickness in several brain regions. **B**: The strongest association was also with left dorsomedial prefrontal cortex area (dmPFC). **C**: The mediation model showed a significant indirect effect, such that lactate+ at dACC partially but significantly mediated the effects of dmPFC on salivary cf-mtDNA response to the psychological stress. a.u.: arbitrary unit.

### Clinical and demographic correlates of mtDNA response to stress

Developmental trauma as reported by CTQ (available in twenty participants) was not significantly associated with salivary cortisol response (r=0.14, p=0.57) but significantly associated with salivary cf-mtDNA response (r=0.50, p=0.031). Recent stress levels as reported by PSS score (N=20) was not significantly associated with either cortisol (r=-0.26, p=0.28) or cf-mtDNA (r=0.04, p=0.87) responses. None of the processing speed (N=33) or the working memory (N=35) tasks were significantly associated with cortisol (r=-0.13 to -0.14, p=0.48 to 0.43) or cf-mtDNA (r=-0.04 to -0.23, p=0.80 to 0.19) response. Age had a U-shape effect on salivary cf-mtDNA response based on a significant quadratic regression (F=4.37, p=0.021) (**Figure 5**); removing the 2 participants who were possible outliers on the age effect (arrows in **Figure 5**), the quadratic effect was even more apparent (R^2^=33.8%, F=7.41, p=0.003). Sex difference on cf-mtDNA response was not significant (p=0.27).

**Figure 5:**
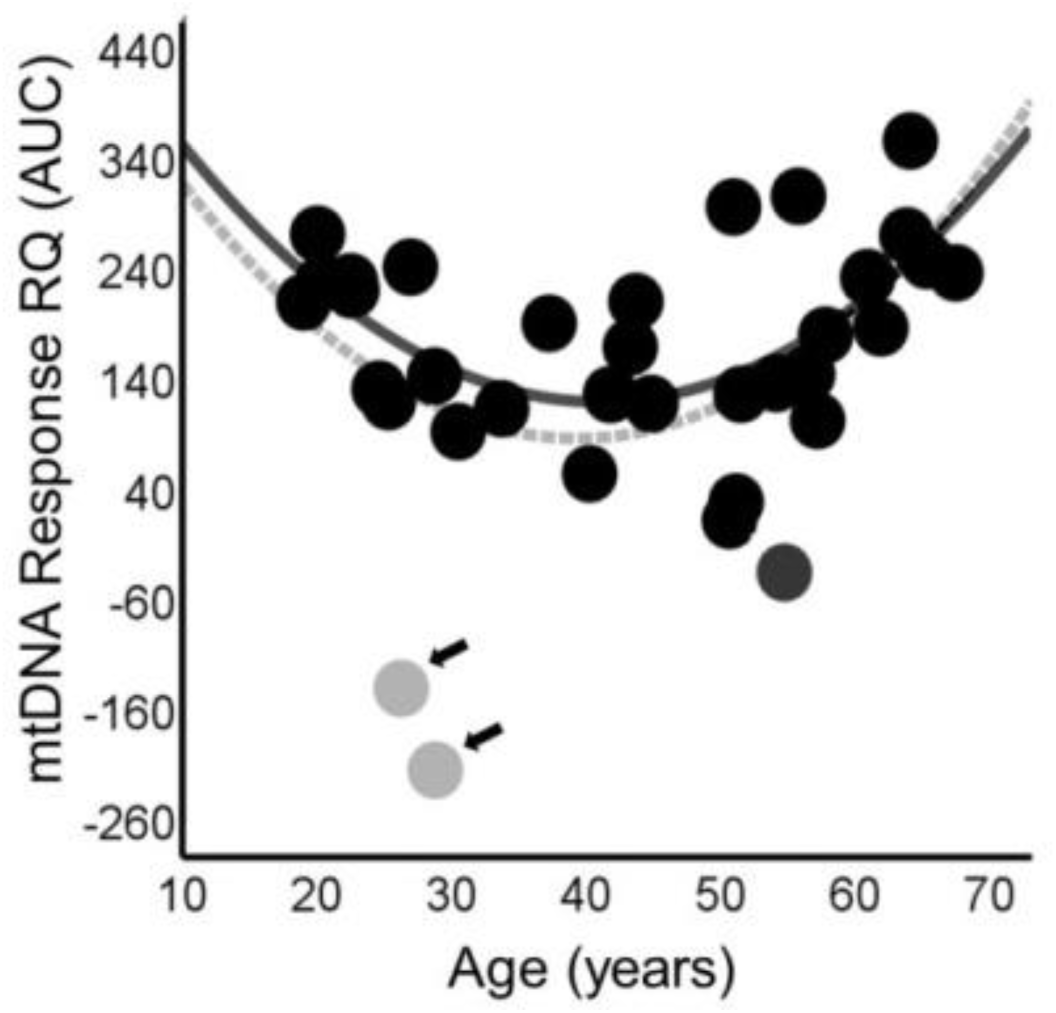
A significant, U-shaped age effect on salivary cf-mtDNA response. This effect was significant in the whole sample (dotted gray line), and the effect was even stronger if the two apparent outliers for the age effect (arrows) were removed (solid black line).

## Discussion

The study found that a brief psychological stress can induce a robust salivary cf-mtDNA release in healthy individuals. This response had only a marginal association with cortisol response. Instead, this stress-induced cf-mtDNA response was strongly associated with the mitochondrion-related metabolite lactate+ measured at the dACC, while thinner cortex especially at the dmPFC was significantly associated with heightened stress-induced cf-mtDNA responses.

The study first attempted to address the apparent proposition that this mitochondrial response was regulated by HPA-led glucocorticoid response. This is because glucocorticoid receptors are abundant in mitochondria [68, 69] and stress has multiple direct and indirect, biphasic effects on mitochondrial integrity [70–72]. Stress response demands high energy supplies, which are normally fulfilled by the highly dynamic mitochondrial fission and fusions and other changes that rapidly respond to energy demands [73, 74]. Excessive glucocorticoids and chronic stress also impair mitochondrial trafficking and induce mitochondrial accumulation and disruptions [4, 72, 75], which may lead to release of mtDNA fragments into the circulation. Despite these clear stress-mitochondrial associations, whether and how a brief psychological stressor may engage this glucocorticoid-mitochondrial response process in humans was previously unclear. We found that there appeared an increase in the correlation between cortisol and cf-mtDNA responses immediately after the stress challenge (**Figure 1E**), but exploratory modeling yielded not statistically significant causal relationship, suggesting that this cf-mtDNA response was unlikely a primarily glucocorticoid-dependent process.

Indeed, mitochondrial response to stress is controlled through multiple routes [1]. For example, acute psychological stress in mice showed that, besides glucocorticoids, mitochondrial responses are also driven or modulated by catecholamines, inflammatory cytokines, whole-body metabolites, and gene expression responses [76]. As discussed earlier, a study using the psychosocial stress test found robust cortisol responses in the saliva while the salivary cf-mtDNA response was not significant [7], which indirectly supports the conclusion of our modeling results that cortisol response may not be the primary driver for the cf-mtDNA response. However, these are cross-sectional analyses; causally designed experiments would be needed to confirm. It is also possible that some of these other pathways may underlie the psychological stress – cf-mtDNA response observed in humans, which warrants additional investigations.

Cf-mtDNA appeared a robust biomarker in rodents for stress-induced mitochondrial disruption. Importantly, inhibition of the Toll-like receptor 9 (TLR-9) can attenuate the cf-mtDNA release and rescue the associated social deficit behaviors, suggesting potentials for developing pharmacological interventions to normalize the stress–mitochondrial cascade [9]. Whether this stress - cf-mtDNA relationship can be assayed through saliva was previously unclear. As discussed in the introduction, TSST induced significant plasma cf-mtDNA release but failed to elicit significant salivary cf-mtDNA response [7]. The plasma mtDNA increase was peaked 45 minutes after TSST and did not trend towards baseline), raising questions on the nature of this delayed cf-mtDNA peak. A more rigorously designed cross-over study in a large sample confirmed that the cf-mtDNA changes observed in serum and plasma were not primarily from the psychosocial stress challenges [13]. Using the psychological stress challenge here, salivary mtDNA responses were peaked immediately after and trended towards baseline after (**Figure 1A**). It is unclear the factors that have contributed to the cf-mtDNA response differences between these two paradigms. Despite some differences, together these data suggest the feasibility to clinically assess mitochondrial response to stress. Cf-mtDNA response to laboratory stressors, especially through saliva sampling, may provide a new way to translate the robust stress–mitochondrial pathway knowledge learned from animal and cellular studies to clinical investigation.

Of its many functions, dACC is strongly associated with stress regulations. Meta-analyses have identified dACC to be the region most strongly associated with fear conditioning among all brain regions, and the superior frontal gyrus is also one of the top regions [18, 77]. These findings have led to the formulation that the medial ‘cingulofrontal cortex’ to be the central autonomic–interoceptive nexus for maintaining homeostasis during heightened arousal states [18]. This network view of the mPFC stress regulation appears supportive of our findings of associations between higher salivary stress response and more lactate+ at dACC and more cortical thinning at dmPFC, but with a novel observation that these associations are linked to salivary mtDNA response to stress.

Our study did not directly examine brain metabolite or brain mitochondrial response to stress, which would require a different study design. Rather, we observed that dACC lactate+ level [26, 78–82] has a negative association with cf-mtDNA response to stress, such that individuals with higher basal lactate+ levels would have stronger salivary cf-mtDNA response, an effect that appeared quite specific compared to other metabolites obtained by ^1^H-MRS (**Figure 2C**). Furthermore, the relationship between cf-mtDNA responses and dmPFC cortical thickness was significantly mediated by dACC lactate+ level (**Figure 4C**). Lactate is thought to link to mitochondrial functioning especially under stressed environment [24, 71, 78, 81]. Besides being a product of anaerobic metabolism, lactate is directly oxidized by mitochondria especially during high energy demand for maintaining brain energy homeostasis [25, 26, 28, 29]. It is unclear how dACC lactate+ levels measured at baseline has such a relatively specific association with stress-induced salivary cf-mtDNA elevation, but the finding is consistent with the notion that lactate accumulation is associated disruptions of mitochondrial functions [83–85] and is indicative of a baseline mitochondrial vulnerability [24, 27].

On brain regions associated with cf-mtDNA vs. cortisol response, the patterns appear quite different. Cortisol response was associated more with cortical thickness at parahippocampal and orbitofrontal regions. In disorders with glucocorticoid dysfunction, key regions affected are the hippocampus and prefrontal cortex [19], which could be viewed as consistent to our cortisol response findings. Interestingly, healthy individuals with stronger cortisol response showed positive associations with cortical thickness in these regions. This is likely because, unlike glucocorticoid increases in chronic stress or glucocorticoid disorders, transient increase of cortisol is physiological and can improve cognitive functioning during stressful situations [86, 87]. The cortisol-related findings contrasted to the primarily mPFC-related associations with cf-mtDNA response. MPFC in rodents or dmPFC in humans plays a key role for stress and emotion regulations [79, 82], potentially explaining why cortical thickness at dmPFC was more strongly associated with salivary cf-mtDNA response to psychological stress. Regardless of the underlying mechanism, a relevant point here is that the patterns of cortical associations between glucocorticoid response vs. cf-mtDNA response to stress appear different (comparing **Figure 3A** vs. **3C**), further suggesting that the cf-mtDNA response may not be primarily glucocorticoid-driven.

The study has several limitations. The brain imaging was not conducted under stress challenges, limiting the interpretations to how these ‘trait’ like brain measures may contribute to the saliva-based cf-mtDNA response. Studies to simultaneously measure brain and saliva response may yield additional insight. We did not examine cf-mtDNA response in psychiatric illnesses, although to first evaluate these hypotheses in healthy people should provide a stronger foundation for future disease research applications. The PRESS sequence with TE=30 ms is not optimized for lactate quantification, which ideally requires a longer TE (e.g., ∼144 ms) or spectral editing techniques to improve specificity, although our study aimed to evaluate a broad range of metabolites [88, 89]. Future studies employing optimized sequences are needed to confirm these findings. Additional studies are also warranted to determine if saliva and plasma cf-mtDNA levels represent same or different cellular origins. Directly assay the mitochondrial morphology, mtDNA gene expression or mutation, and other mitochondrial proteins were not performed in this study but may inform the psychological stressor – cf-mtDNA response mechanisms and should be considered in future studies. Finally, the study used only small to modest samples, and therefore these findings should be interpreted cautiously and require replication in larger sample studies.

In summary, this paradigm offers a potentially new, non-invasive approach to evaluate the stress-induced mitochondrial functioning in human subjects. It uses simple saliva collection through a well-established psychological stress paradigm and cf-mtDNA assay. The evidence so far suggests that this cf-mtDNA response is not primarily glucocorticoid-dependent but rather associated with brain medial prefrontal metabolite and structural markers thought to be related mitochondrial function and the central stress regulation hub, respectively. Further underlying pathway investigations are necessary to determine the value of this approach in studying the stress-mitochondrial pathways in health, aging, psychopharmacology, and neuropsychiatric conditions where psychological stress plays a role.

## Acknowledgements

Funding support was received from NIH grants R01MH133812, R01MH116948, R01MH112180 (LEH), MH120876 and MH128771 (AP). The findings and conclusions in this article are those of the authors and do not necessarily represent the views or opinions of these organizations, which had no role in designing or planning this study; in collecting, analyzing, or interpreting the data; in writing the article; or in deciding to submit this article for publication.

